# Comprehensive analysis of end-modified long dsDNA donors in CRISPR-mediated endogenous tagging

**DOI:** 10.1101/2024.06.28.601124

**Authors:** Rioka Takagi, Shoji Hata, Chiharu Tei, Akira Mabuchi, Ryosuke Anzai, Masamitsu Fukuyama, Shohei Yamamoto, Takumi Chinen, Atsushi Toyoda, Daiju Kitagawa

## Abstract

CRISPR-mediated endogenous tagging is a powerful gene editing technique for studying protein dynamics and function in their native cellular environment. While the use of 5’ modified DNA donors has emerged as a promising strategy to improve the typically low efficiency of knock-in gene editing, the underlying mechanisms remain poorly understood. In this study, we conducted a comprehensive analysis of end-modified long linear dsDNA donors in CRISPR-mediated endogenous tagging in human non-transformed cells. In-depth analysis of repair patterns reveals that 5’ biotinylation of dsDNA donors significantly reduces imprecise insertions, thereby enhancing homology-directed repair (HDR)-mediated precise insertion efficiency. Notably, the impact of biotinylation on repair patterns resembles that of non-homologous end joining (NHEJ) pathway inhibition, suggesting its role in preventing NHEJ-mediated mis-integration. Moreover, combining biotin modification with NHEJ inhibitor treatment further improves bi-allelic knock-in efficiency. Overall, this study provides novel insights into the mechanisms by which 5’ modifications enhance precise knock-ins and demonstrates their potential for achieving high-efficient, prercise endogenous tagging in human cells.

## Introduction

CRISPR-mediated endogenous tagging is a powerful genome editing technique that utilizes the CRISPR/Cas system to precisely insert a desired a tag sequence, such as a fluorescent protein, affinity tag, or enzyme, into the endogenous locus of a gene of interest (Hsu et al., 2014). This approach enables researchers to study the expression, localization, and function of proteins in their natural cellular context. This technique involves designing a guide RNA (gRNA) that targets the desired insertion site, preparing a donor DNA template containing the sequence to be inserted flanked by homology arms (HA), and delivering the CRISPR components including the gRNA, Cas nuclease, and donor DNA into the target cells. Cas9 and Cpf1 are commonly used Cas proteins for knock-in experiments, recognizing different PAM sequences and producing different DNA end structures after cleaving the target site (Gasiunas et al., 2012; Zetsche et al., 2015). Guided by the gRNA, the Cas nuclease creates a double-strand break (DSB) at the target site (Jinek et al., 2012), which is then repaired by DNA repair mechanisms, such as homology-directed repair (HDR), using the donor DNA as a template (Yao et al., 2018). This technique has revolutionized genome editing and has been applied in various fields of basic research. However, significant challenges remain to be addressed, notably low knock-in efficiency and limited accuracy, particularly in non-transformed cells.

While the HDR pathway enables the precise incorporation of the donor sequence flanked by HAs into the target site during repair, the Cas-induced DNA break can also be repaired by other DNA repair pathways in a homology-independent manner. Non-homologous end joining (NHEJ) is the most dominant non-HDR pathway, re-ligating exposed DNA ends resulting from the break (Chang et al., 2017a; Decottignies, 2013). NHEJ is often error-prone, resulting in insertions or deletions (indels) that can disrupt gene function. Moreover, NHEJ is implicated in joining the break ends with donor DNA ends, which causes imprecise donor integrations (Denes et al., 2021). Other non-HDR pathways include microhomology-mediated end joining (MMEJ) and single-strand annealing (SSA), which uses short and long homologous sequences respectively for repair and can cause deletions and chromosomal rearrangements (Bhargava et al., 2016; Ceccaldi et al., 2016; Scully et al., 2019; Seol et al., 2018; Sfeir and Symington, 2015; Sinha et al., 2016). These pathways compete with HDR for DSB repair and can reduce the efficiency and precision of CRISPR-mediated knock-in, thereby prompting the development of strategies to suppress them and improve efficiency of precise genome editing outcomes (Fu et al., 2021; Maruyama et al., 2015; Schimmel et al., 2023; Tei et al.; van de Kooij et al., 2022; Yu et al., 2015).

The type of DNA donor used as HDR templates is also a critical determinant of the efficiency and accuracy of gene editing. Single-strand DNA (ssDNA) is preferred to insert short sequences (Chen et al., 2011; Wu et al., 2013; Zhang et al., 2022). For inserting longer sequences such as fluorescent protein tags, linear double-strand DNA (dsDNA) donors have been shown to provide higher knock-in efficiency compared to ssDNA donors and circular plasmid DNA donors, a classical DNA donor type (Canaj et al., 2019; Mabuchi et al., 2023). However, regardless of the donor type, various inaccurate incorporations of donor sequences into the target sites can occur, such as blunt insertion (the end of the donor DNA, including the HAs, are directly ligated to the cut site), imperfect insertion (the end of the donor DNA is trimmed and then incorporated) and concatenation (multiple copies of the donor DNA join end-to-end to form a tandem array), rather than solely precise insertions referred to as perfect HDR.

5’ end modification of linear DNA donors, such as the incorporation of a biotin molecule to the 5’ end of a DNA donor, has emerged as a promising strategy to enhance the efficiency of knock-in gene editing. Studies have reported significant improvements in knock-in efficiency, with up to a several-fold increase compared to unmodified donors, in various cell types (Canaj et al., 2019; Ghanta et al., 2021; Gutierrez et al., 2018; Yu et al., 2020). As a simple and cost-effective approach, 5’ modification of DNA donors has the potential to become a standard practice in gene knock-ins. However, the mechanisms underlying enhanced efficiency of endogenous tagging with 5’ modified long linear dsDNA donors remain elusive, primarily due to the lack of evidence regarding the influence of such modifications on repair patterns in these donors.

In this study, we performed comprehensive analysis of end-modified long linear dsDNA donors in CRISPR-mediated endogenous tagging in human diploid non-transformed cells. Our results revealed that 5’ biotinylation enhanced overall donor integration while reducing the proportion of imprecise insertion, thereby enhancing efficiency of HDR-mediated precise insertion. This impact of biotinylation on repair patterns resembled that of NHEJ inhibition, suggesting its role in preventing NHEJ-mediated mis-integration. Moreover, combining biotin modification with NHEJ inhibitor treatment further improved bi-allelic knock-in efficiency. Therefore, this study provides novel insights into the mechanisms by which 5’ modification enhances the efficiency of precise knock-in, and propose that combining this with NHEJ inhibition as a strategy for achieving high-efficient, precise endogenous tagging in human cells.

## Results

### 5’ modifications of dsDNA donor enhance the efficiency and accuracy of CRISPR knock-in with long DNA sequences

Among several end modifications of linear dsDNA donors, 5’ biotinylation and AmC6 modification are known to increase CRISPR-mediated knock-in efficiency (Gutierrez et al., 2018; Yu et al., 2020). Therefore, we first examined the effects of these end modifications on knock-in outcomes in the hTERT-immortalized RPE1 cell line, a human non-transformed diploid cell line commonly used in cell biology. Knock-in experiments in RPE1 cells were conducted using a modified version of the previously established cloning-free endogenous tagging method (Mabuchi et al., 2023) (Fig. 1a). Linear donor DNA containing 90 bases of homology arm (HA) sequences was prepared by PCR using primers with or without 5’ end modifications. Recombinant Cas nucleases and *in vitro* transcribed guide RNAs were mixed to form ribonucleoprotein (RNP) complexes, which were then electroporated into cells along with the donor DNA (Fig. 1a).

**Fig. 1:**
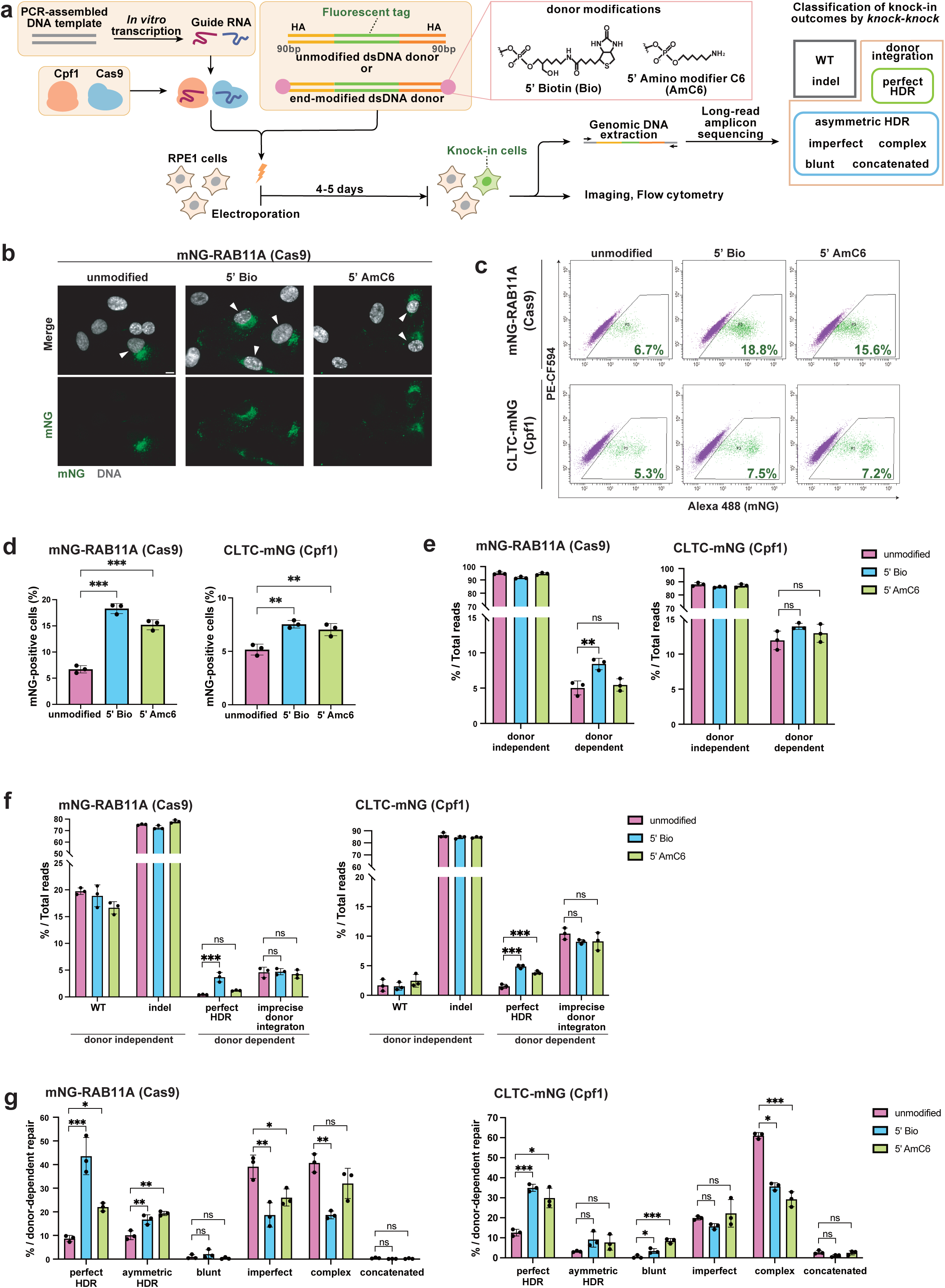
5’ modifications of dsDNA donor enhance the efficiency and accuracy of long-sequence knock-in using CRISPR/Cas system. **a**, Schematic overview of the experimental system for evaluating endogenous gene tagging in human RPE1 cells. Donor DNA was PCR-amplified as shown in material and methods and in Supplementary Table 1. Guide RNA transcribed *in vitro* from PCR-assembled DNA was mixed with commercially available Cpf1 or Cas9 proteins to form RNP complexes, and these RNPs and the dsDNA donors were electroporated into RPE1 cells. The fluorescent protein sequence is expected to be inserted into the target locus of Cas nucleases by using the donor DNA as a HDR template. The electroporated cells were subsequently cultured in a medium for 4-5 days and then subjected to fluorescence imaging, flow cytometric analysis, or genome extraction. Extracted genome was PCR-amplified for long-read amplicon sequencing by PacBio and subsequent *knock-knock* analysis of knock-in outcomes. **b**, Representative images from cells with Cas9-mediated mNG tagging of RAB11A. Arrowheads indicate cells with endosomal mNG signals. Cells were fixed and analyzed 5 days after electroporation. Scale bar: 10 µm. **c**, Flow cytometric analysis of Cas9-mediated mNG-RAB11A and Cpf1-mediated CLTC-mNG knock-in cells. Cells were analyzed 4 days after electroporation. Percentages of cells with mNG signal are shown in the plots. **d**, Quantification of percentages of mNG-positive cells from (**c**). Data from three biological replicates are shown. Approximately 10,000 cells were analyzed for each sample of RAB11A and CLTC. **e**, Distribution of donor-independent or donor-dependent repair across the targeted gene loci. 10,873-50,357 reads were analyzed for each sample. **f**, A brief classification of donor-independent repair and donor-dependent repair across the targeted gene loci from (e). **g**, Percentage of perfect HDR and imprecise integration events within donor-dependent repair for the targeted genes. For each sample, 1,473-6,189 reads categorized as the donor-dependent repair were analyzed. In **d**, **f**, and **g**, data from three biological replicates are represented as mean ± S.D. and P value was calculated by a Tukey–Kramer test. ***P < 0.001, **P < 0.01, *P<0.05, ns: Not significant.

To investigate the effects of dsDNA donor end modifications on both Cas9- and Cpf1-mediated knock-in, we performed Cas9-mediated N-terminal tagging of the trafficking protein RAB11A and Cpf1-mediated C-terminal tagging of the vesicle coating protein CLTC using the green fluorescent protein mNeonGreen (mNG) (Fig. 1a). Fluorescence imaging revealed that most mNG-positive cells exhibited the expected localization corresponding to each mNG-fused endogenous protein. Consistent with previous reports (Yu et al., 2020), 5’ modifications increased the proportion of cells with mNG signal (Fig. 1b). To precisely quantify knock-in efficiency, we performed flow cytometric analysis 4 days after electroporation. This analysis showed that dsDNA donor modified with 5’ biotin (labeled as 5’ Bio in the Figure) or 5’ AmC6 increased Cas9-mediated knock-in efficiency at the *RAB11A* locus by approximately 2.7-fold and 2.3-fold (from 6.7% to 18.3% and 15.2%), respectively, compared to the unmodified control (Fig. 1c, d). On the other hand, the increase in Cpf1-mediated knock-in efficiency at the *CLTC* locus was limited to approximately 1.5-fold for both modifications (from 5.2% to 7.5% and 7.0%).

After confirming that the 5’ modifications of long dsDNA donor significantly enhance knock-in efficiency, we further verified the resulting repair patterns by performing long-read amplicon sequencing of the knock-in alleles using PacBio and genotyping using a computational framework called *knock-knock* (Canaj et al., 2019; Tei et al., 2023). Following electroporation with the different donors, the knock-in target sites were amplified by PCR from the extracted genomic DNA (Fig. 1a). During the subsequent sequencing process, each Hi-Fi read was classified through the *knock-knock* classification process into specific DSB repair outcomes: WT or indels (donor-independent repair), or perfect HDR or inaccurate integration subtypes (donor-dependent repair). Interestingly, 5’ biotinylation, but not 5’ AmC6 modification, increased donor-dependent repair in the Cas9-mediated knock-in at the *RAB11A* locus (Fig. 1e). A similar tendency was also observed for the Cpf1-mediated knock-in at the *CLTC* locus. Among donor-dependent integrations at the *RAB11A* locus, 5’ biotinylation significantly increased the proportion of perfect HDR, while 5’ AmC6 modification had a mild effect (Fig. 1f). In contrast, at the *CLTC* locus, both modifications significantly increased perfect HDR. Regarding donor-independent repair, 5’ modifications did not impact the repair patterns overall, except for the case of AmC6 at the *RAB11A* locus (Fig. S1b).

Based on the classification by *knock-knock*, we categorized imprecise integration events into the following: asymmetric HDR (only one side of the donor DNA is precisely integrated in an HDR manner), blunt (both ends of the donor DNA, including the HAs, are directly ligated to the cut site), imperfect (at least one end of the donor DNA is trimmed and incorporated), concatenated (multiple insertions of donors), and complex (not classified into the other four mis-integration categories). In the donor-dependent repair detected in Cas9-based endogenous tagging at the *RAB11A* locus, 5’ biotinylation significantly decreased the proportion of HDR-independent repairs, such as imperfect (from 39.1% to 18.7%) and complex (from 40.7% to 18.7 %), and instead markedly increased the proportion of perfect HDR (from 8.7% to 43.6%) and slightly increased that of asymmetric HDR (from 10.1% to 16.7%) by approximately 5-fold and 1.6-fold, respectively (Fig. 1g). The 5’ AmC6 modification also altered the repair pattern similarly, albeit less dramatically. These data suggest that 5’ modifications of linear long dsDNA donor shift repair patterns from non-HDR to HDR ones, with the magnitude of this effect varying by the modification type.

Unlike at the *RAB11A* locus, the impact of both 5’ modifications on donor-dependent repair patterns was similar at the *CLTC* locus (Fig. 1g). Specifically, both modifications significantly increased the proportion of perfect HDR and also tended to increase asymmetric HDR in donor-dependent repairs. Interestingly, while both modifications markedly reduced the proportion of complex repairs, they did not exhibit a pronounced inhibitory effect on imperfect repairs, in contrast to the *RAB11A* locus. Overall, these data suggest that 5’ modifications of long linear dsDNA donor positively affect the accuracy of knock-in, likely by suppressing imprecise non-HDR repairs. However, the extent of this effect varies depending on the type of 5’ modification and the Cas system used. Our analyses demonstrate that 5’ biotinylation dramatically enhances the efficiency of accurate knock-ins in both Cas9 and Cpf1 systems.

### Doggybone DNA donors enhance donor-dependent repair but not its precise integration

To investigate the effect of another type of DNA end protection in CRISPR-mediated knock-in with long linear dsDNA donors, we next focused on doggybone DNA (dbDNA). dbDNA is a unique DNA structure in which both ends of a linear double-stranded DNA are connected by hairpin loops (Fig. 2a; Deneke et al., 2002; Heinrich et al., 2002), and it has been shown to be useful in various applications, serving as an alternative to traditional plasmid vectors for producing iPSCs (Thornton et al., 2021), lentiviruses (Karda et al., 2019), and CAR T cells (Bishop et al., 2020), and also in DNA vaccines (Allen et al., 2018). The structure of dbDNA effectively protects the DNA ends and enhances their stability (Deneke et al., 2002), making it a potentially attractive option for improving the efficiency and accuracy of CRISPR-mediated knock-in. To generate dbDNA donor, we reacted recombinant TelN protelomerase with a plasmid containing mNG coding sequence flanked by 90 bp HAs and 56 bp TelRL sequences (Heinrich et al., 2002). TelN protelomerase cut the plasmid at the middle of the TelRL sequence and leaves covalently closed ends at the site of cleavage, generating two different dbDNAs. To purify the desired dbDNA donor, the other dbDNA containing plasmid backbone sequence was digested by exonucleases following treatment with restriction enzymes (Fig. 2a, b).

**Fig. 2:**
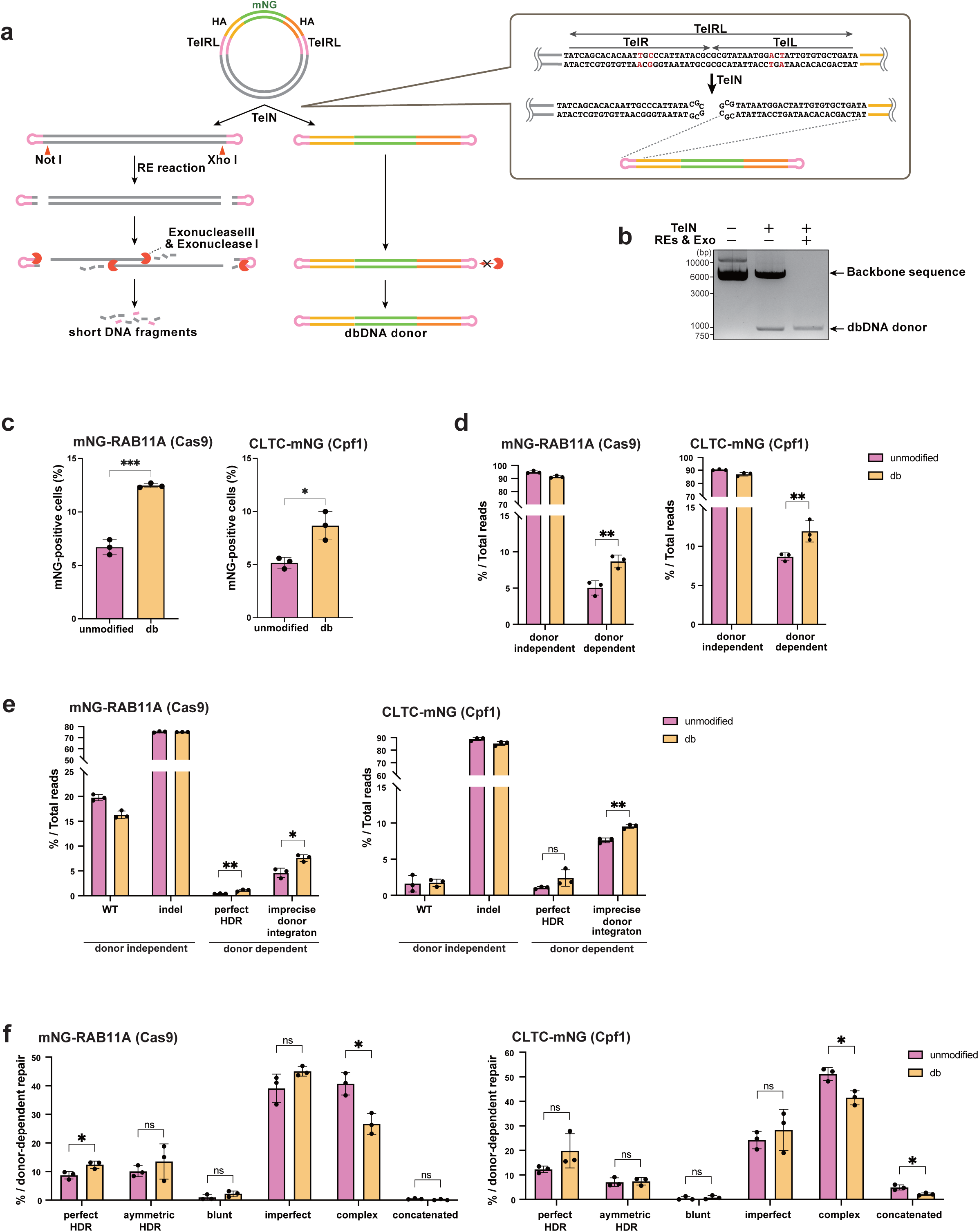
Doggybone DNA donors improve donor-dependent repair but not its precise integration. **a**, Schematic overview of doggybone DNA preparation process. A template plasmid is designed to contain mNG coding sequence flanked by 90 bp HAs and 56 bp TelRL sequences. The TelRL sequence is shown in the right. After TelN protelomerase reaction, a 28bp TelL sequence remains at both ends of the homology arm, and the backbone DNA with closed ends is digested by restriction enzymes followed by exonucleases, allowing for the purification of only dbDNA. **b**, Confirmation of doggybone DNA production by the method shown in (**a**). The plasmid template of the donor DNA for Cas9-mediated mNG tagging of RAB11A was used in this Figure. **c**, Percentages of mNG-positive cells quantified by flow cytometric analysis in Cas9-mediated mNG-RAB11A and Cpf1-mediated CLTC-mNG knock-in cells. After electroporation, the cells were cultured for 4 days before the analysis. Data from three biological replicates are shown. Approximately 10,000 cells were analyzed for each sample. **d,** Distribution of donor-independent or donor dependent repair across the targeted gene loci. 9,198-47,322 reads were analyzed for each sample. **e**, A brief classification of donor-independent repair and donor-dependent repair from (d). Data are represented as mean ± S.D. **f**, Percentage of perfect HDR and imprecise integration within donor-dependent repair across the targeted genes. For each sample, 1,133-36,115 reads categorized as the donor-dependent repair were analyzed. In **c**, **d**, **e**, and **f**, data from three biological replicates are represented as mean ± S.D. A two-tailed, unpaired Student’s t test was used to obtain the P-value. ***P < 0.001, **P < 0.01, *P<0.05, ns: Not significant.

Using dbDNA donors, we performed Cas9 and Cpf1-mediated endogenous tagging of mNG to the *RAB11A* and *CLTC* alleles, respectively. FACS analyses showed that dbDNA donors significantly increased both Cas9 and Cpf1-mediated knock-in efficiencies (Fig. 2c). We next assessed the repair patterns in endogenous tagging with dbDNA donors. The long-read amplicon sequencing and subsequent *knock-knock* analysis revealed that the dbDNA donor resulted in a significant increase in donor-dependent repair, similar to the case of 5’ biotinylation (Fig. 2d). Given that dbDNA exhibits high stability due to end protection, the increase in donor-dependent repair likely reflects the enhanced intracellular stability of the donor. However, to our surprise, dbDNA donors did not show a marked increase in perfect HDR, unlike the case with 5’ biotinylation (Fig. 2e). The proportion in donor dependent repair was only slightly different between unmodified and dbDNA donors, except for complex repair (Fig. 2f). It has been suggested that even when fluorescent proteins are inaccurately knocked-in, they may retain their fluorescence properties(Feng et al., 2017). Therefore, the increment of donor-dependent integration, including imprecise knock-in, may contribute to a higher number of mNG positive cells with dbDNA donors in endogenous tagging. However, in terms of the accuracy of knock-in, dbDNA donor is likely not a better option compared to 5’ biotinylation.

### 5’ biotinylation suppresses NHEJ-mediated inaccurate repair with long linear double-stranded DNA donors

Next, we investigated why 5’ modification increases the accuracy of knock-in. To explore the possibility that 5’ modification indirectly increases perfect HDR outcomes by suppressing non-HDR, we analyzed impact of the inhibition of NHEJ, the major non-HDR pathway, on CRISPR-mediated endogenous tagging with 5’ modified linear dsDNA donors. As reported previously (Fu et al., 2021; Maruyama et al., 2015), treating cells with an NHEJ inhibitor dramatically increased knock-in efficiency with unmodified dsDNA donor in Cas9- and Cpf1-mediated endogenous tagging at the *RAB11A* and *CLTC* loci by approximately 2.4-fold and 2-fold (from 6.7% to 16.1% and from 5.2% to 11.6%), respectively (Fig. 3a, b). Interestingly, however, the combination of 5’ modification with NHEJ inhibitor did not provide further enhancement of knock-in efficiency at either locus compared to sole inhibition of NHEJ or even reduced the efficiency in the case of the *RAB11A* locus. This suggests that the improvement of knock-in efficiency by the 5’ modifications at least partially depend on the suppression of NHEJ-mediated knock-in mechanisms.

**Fig. 3:**
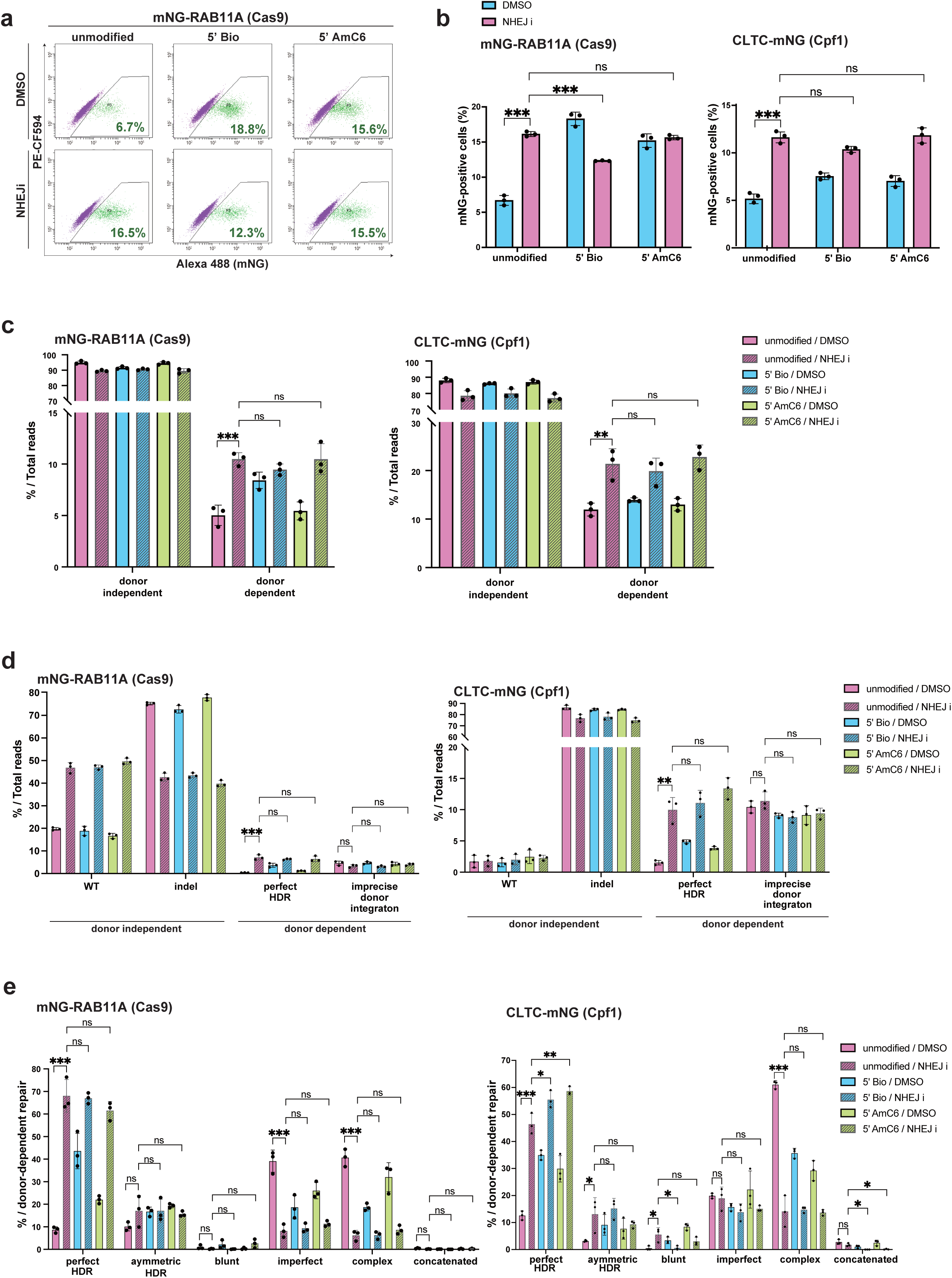
5’ biotinylated donor partially inhibits NHEJ-dependent integration and enhances HDR efficiency. **a**, Flow cytometric analysis of Cas9-mediated mNG-RAB11A knock-in cells with or without NHEJi treatment. After electroporation, cells were cultured for 24 hours in medium containing either DMSO or 1 µM NHEJ inhibitor, followed by an additional 3-day culture in fresh medium before conducting flow cytometric analysis. Percentages of cells with mNG signal are shown in the plots. **b**, Quantification of percentages of mNG-positive cells from (a) and Cpf1-mediated CLTC-mNG knock-in cells. Data from three biological replicates are shown. Approximately 10,000 cells were analyzed for each sample of RAB11A and CLTC. **c**, Distribution of donor-independent or donor dependent repair across the targeted gene loci. 9,230-50,357 reads were analyzed for each sample. **d**, A brief classification of donor-independent repair and donor-dependent repair from (c). **e**, Percentage of perfect HDR and imprecise integration within donor-dependent repair across the targeted genes. For each sample, 1,473-10,419 reads categorized as the donor-dependent repair events were analyzed. In **a-e**, all data treated with DMSO are identical to those in Fig. 1. Since they were obtained in the same experiment as the data for NHEJi treatment, a comparative analysis is feasible. Data are represented as mean ± S.D. and P value was calculated by a Tukey– Kramer test. ***P < 0.001, **P < 0.01, *P<0.05, ns: Not significant.

NHEJ inhibition significantly altered repair outcomes at both loci, reducing donor-independent repairs while increasing donor-dependent repairs, particularly perfect HDR (Fig. 3c, d). At the *RAB11A* locus, where Cas9-based endogenous tagging was applied, NHEJ inhibition markedly decreased the proportion of HDR-independent repairs: imperfect repairs decreased from 39.1% to 8.2% and complex repairs from 40.6% to 6.2% in donor dependent repairs (Fig. 3e). Instead, it significantly increased the proportion of perfect HDR (from 8.7% to 68.0%) and slightly increased asymmetric HDR (from 10.1% to 17.1%) by approximately 8-fold and 1.7-fold, respectively. Similarly, at the *CLTC* locus, where Cpf1-based endogenous tagging was applied, NHEJ inhibition dramatically increased HDR-mediated repairs (from 12.6% to 46.5% for perfect HDR; from 3.2% to 13.1% for asymmetric HDR), although its effects on non-HDR patterns differed from those at the *RAB11A* locus, with imperfect repair remaining unaffected (from 19.9% to 19.0%) (Fig. 3e). Notably, the biases in donor-dependent repair patterns caused by NHEJ inhibition closely resemble those of 5’ modification. Furthermore, combining 5’ modification with NHEJ inhibition did not provide additional enhancement of HDR-mediated repair or further reduction of non-HDR at either locus compared to NHEJ inhibition alone, mirroring the results observed for knock-in efficiency. The similarity in the biases of donor-dependent repair patterns between 5’ modification and NHEJ inhibition, coupled with the absence of additive effects, suggests that 5’ modification may inhibit donor-dependent imprecise repair mediated by NHEJ.

### The combination of 5’ biotinylated donor DNA and NHEJ inhibitor treatment boosts bi-allelic knock-in efficiency

When establishing endogenously tagged cell lines with fluorescent proteins, flow cytometric cell sorting is commonly employed to enrich cells with successful knock-in. To investigate the effects of 5’ biotin, NHEJ inhibition or the combination of both in these practical scenarios, we collected mNG-positive cells using flow cytometry after endogenous mNG tagging at the *RAB11A* locus and then performed long-read amplicon sequencing and *knock-knock* analysis on these enriched cell populations. Surprisingly, unlike the results obtained without sorting, the combination of 5’ biotinylation of donor DNA and NHEJ inhibition resulted in an additive increment of donor dependent integration, especially perfect HDR, in the mNG-sorted cells (Fig. 4a, b). The increase in perfect HDR in the combination condition was 8-fold (from 5.7% to 46.2%) compared to the control condition with unmodified donor and without NHEJ inhibition. Interestingly, however, the combination of 5’ modification with NHEJ inhibitor did not provide further enhancement of the proportion of perfect HDR within donor-dependent repairs compared to NHEJ inhibition alone (Fig. 4c). Besides the proportion of perfect HDR, 5’ modification had no impact on donor-dependent repair patterns under NHEJ inhibition in sorted cells, which is similar to the results observed in non-sorted cells (Fig. 3e). These data indicate that, within the enriched population of edited cells through cell soring, the increase in perfect HDR with 5’ modification under NHEJ inhibition is not due to a reduction in imprecise donor-dependent repair, but to an increase in the overall number of donor-dependent repairs.

**Fig. 4:**
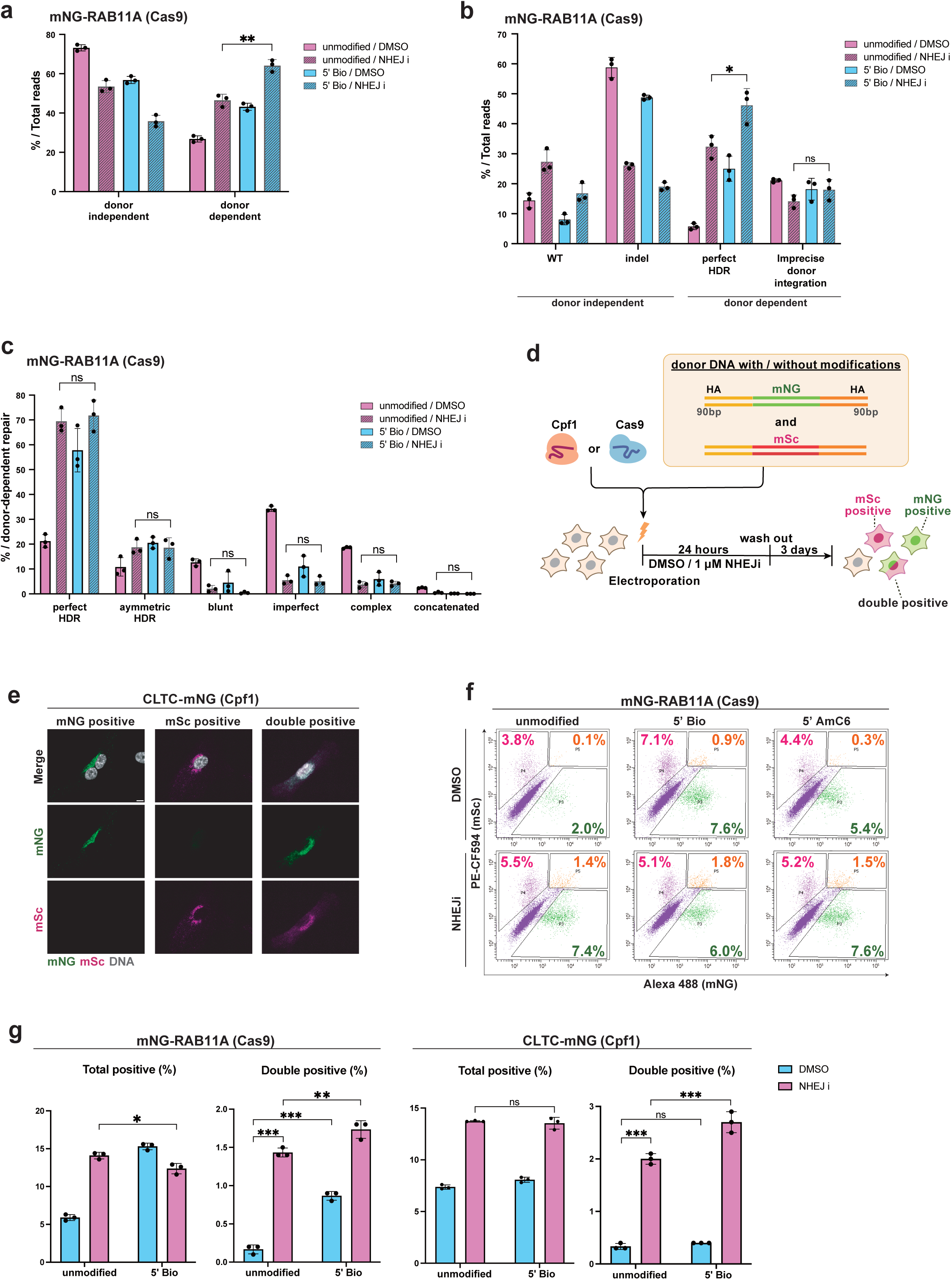
Combination of 5’ biotinylation of donor DNA and NHEJi treatment enhances bi-allelic knock-in. **a**, Distribution of donor independent or donor dependent repair across the targeted gene loci in cells sorted by mNG signal 4 days after electroporation of Cas9-RNP and donor DNA for mNG tagging of RAB11A. 11,796-44,395 reads were analyzed for each sample. **b**, A brief classification of donor-independent repair and donor-dependent repair from (a). **c**, Percentage of perfect HDR and imprecise integration within donor-dependent repair across the targeted genes. For each sample, 7,186-28,760 reads categorized as the donor-dependent repair were analyzed. In **a-c**, data from three biological are represented as mean ± S.D. A two-tailed, unpaired Student’s t test was used to obtain the P-value. **P < 0.01, *P<0.05, ns: Not significant. **d**, Schematic overview of dual-color fluorescent tagging. The same amount of the donors containing either mNG coding sequence or mScarlet (mSc) coding sequence were simultaneously electroplated into cells with Cas-RNPs. **e**, Representative images from cells with Cas9-mediated single- or dual-color tagging of CLTC. Cells at 5 days after electroporation were fixed and analyzed. Scale bar: 10 µm. **f**, Flow cytometric analysis of Cas9-mediated dual-color tagging of mNG and mSc on RAB11A allele of RPE1 cells. Cells at 4 days after electroporation were analyzed. The types of modifications and condition of drug treatment are indicated above and to the left of the plots, respectively. Percentages of cells with mNG, mSc, and both signals are shown in the plots in green, magenta, and orange text, respectively. **g**, Quantification of percentages of cells positive for either mNG or mSc or both (Total positive), or cells positive for both mNG and mSc (Double positive) from (**e**). Data from three biological replicates are shown. Approximately 20,000 cells were analyzed for each sample of RAB11A and CLTC. Data are represented as mean ± S.D. P value was calculated by a Tukey–Kramer test. ***P<0.001, **P < 0.01, ns: Not significant.

Given that the fluorescent cells possess at least one allele repaired with the donor coding mNG sequence, the increase in donor-dependent repairs within these cell populations may result from knock-ins at both copies of the gene per single cell. As the RPE1 cells used in this study are a normal diploid cell line, it is plausible that 5’ biotinylation enhances bi-allelic knock-in efficiency under NHEJ inhibition. To address this possibility, we performed simultaneous dual-color tagging with both mNG and the red fluorescent protein mScarlet (mSc) (Fig. 4d, e). Cas9-mediated dual-color tagging of *RAB11A* resulted in a comparable percentage of total fluorescence-positive cells as single-color tagging of the same gene in all the conditions tested (Fig. 4f, g). Inhibition of NHEJ or 5’ biotinylation increased the percentage of mNG and mSc double-positive cells (from 0.17% to 1.4% and 0.87%, respecively) compared to the control condition. Notably, 5’ biotinylation further increases the double-positive cells under NHEJ inhibition (to 1.7%). The increase in the double-positive cells was approximately 10-fold compared to the control condition with unmodified donor and without NHEJ inhibition. Similarly to the *RAB11A* locus, this combination also increased the proportion of double-positive cells in Cpf1-mediated dual-color tagging of *CLTC*. These data suggest that 5’ biotinylation further enhances the improvement of bi-allelic knock-ins achieved by NHEJ inhibition.

## Discussion

In this study, we comprehensively evaluated the influence of DNA end modifications in CRISPR-mediated endogenous tagging with long linear dsDNA donors. Among the various modifications tested, 5’ biotinylation demonstrated remarkable improvements in efficiency and accuracy of knock-in events (Fig. 1c-g). In-depth analysis of repair patterns at the target site revealed that 5’ biotinylation likely prevents NHEJ-mediated imprecise donor integration, thereby enhancing the efficacy of precise gene knock-in (Fig. 3a-e). Notably, 5’ biotinylation further enhanced the improvement of bi-allelic knock-ins under NHEJ inhibition (Fig. 4a-c, f, g). These enhancements were observed across both Cas9 and Cpf1 systems, highlighting the broad applicability of this modification strategy for high-efficiency, precise endogenous tagging in human cells.

Biotinylation of donor DNA enhanced knock-in efficiency, likely through a combination of two mechanisms: an increase in frequency of total donor integration and an increase in accuracy of integration (Fig. 1e-g). The increase in donor-dependent repair with biotinylation suggests an improvement in donor stability, as similar effects are seen with dbDNA (Fig. 2d), which exhibits high intercellular stability. The high stability of dbDNA is attributed to its closed ends, conferring resistance to exonucleases. Therefore, 5’ biotinylation may similarly provide DNA donors with resistance to 5’ exonucleases, thereby increasing donor-dependent repair. Exonucleases are known to digest polynucleotides from the ends by cleaving phosphodiester bonds. Therefore, it is unlikely that 5’ biotin inhibits exonuclease activity merely by occupying the 5’ DNA end (Lovett, 2011; Sayers and Ecksteins., 1990). Given that 5’ biotin associated with streptavidin inhibits activity of some 5’ exonucleases in vitro (Chang et al., 2015; Cannavo and Cejka., 2014), it is possible that intracellular molecules binding to biotin are involved in this inhibition. Nonetheless, how 5’ biotinylation promotes the stabilization of DNA within cells needs to be elucidated in future studies.

The increased knock-in accuracy observed with 5’ biotinylated donors likely results from the suppression of NHEJ-mediated donor mis-integration, specifically imperfect and complex repairs (Fig. 3e). DSB repairs though NHEJ can follow two paths: one where blunt ends at DSBs are ligated directly to restore original sequence (WT repair), and another where break ends undergo processing before annealing and ligation, leading to indel repair (Chang et al., 2017). The latter pathway involves Artemis nuclease, which can mediate 5’ end resection to convert blunt ends to 3’ staggered ends (Chang et al., 2015). The NHEJ inhibitor used in this study reduced indel formation while increasing WT repair among donor-independent repairs, suggesting it inhibits the processing pathway rather than the non-processing (direct ligation) pathway (Fig. 3d). Interestingly, the effects of the NHEJ inhibitor and donor biotinylation on imperfect and complex repairs differed between Cas9- and Cpf1-mediated knock-ins at the *RAB11A* and *CLTC* loci, respectively (Fig. 3e). While NHEJ inhibition or biotinylation reduced imperfect repairs in Cas9-mediated editing, they did not affect this repair pattern in Cpf1-mediated one. This difference likely stems from the distinct DNA end structures generated by Cas9 and Cpf1 (Jinek et al.; Zetsche et al., 2015).

Cas9 induces double-strand break with blunt ends. Therefore, the dependence on the NHEJ processing pathway of imperfect repair in Cas9-mediated knock-in suggests that this repair process involves processing of the blunt ends at DSBs and the termini of donor DNA to form 3’ staggered ends. As mentioned earlier, if 5’ biotin confers resistance to 5’ exonucleases, it would likely inhibit the formation of 3’ staggered ends on the donor by Artemis, which is known to possess 5’ exonuclease activity to DNA blunt ends. This explanation aligns with the observed reduction of imperfect repairs in Cas9 systems with 5’ biotinylation. In contrast, Cpf1 generates cut ends with 5’ overhangs. These 5’ staggered DSB ends are known to ligate with donors possessing similar 5’ staggered ends through MMEJ (Zhao et al., 2022). Therefore, it makes sense that NHEJ inhibitors or 5’ biotinylation, which potentially inhibit the formation of 3’ staggered ends of donor DNAs, would not impact imperfect repair in Cpf1-mediated knock-ins. Complex repair, characterized by intricate integration of multiple donors, was reduced in both Cas9 and Cpf1 systems through NHEJ inhibition or 5’ biotinylation of the DNA donor. Summarizing the above discussion, the integration of donors in complex repair is likely mediated by the NHEJ processing pathway, possibly involving the formation of 3’ staggered ends on the donor DNA.

Our dual-color endogenous tagging analysis revealed that 5’ biotinylation further enhanced the improvement of bi-allelic knock-ins achieved by NHEJ inhibition (Fig. 4g). This observation aligns with the increased donor-dependent repair patterns observed in fluorescence-sorted cells, which possess at least one fluorescent mNG-edited allele, when 5’ biotinylated donor was used under NHEJ inhibition (Fig. 4a). Interestingly, however, 5’ biotinylation did not affect the proportion of donor-dependent integration without sorting (Fig. 3c), suggesting that 5’ biotinylation increases bi-allelic integration without enhancing overall donor integration. This discrepancy implies a puzzling effect of 5’ biotinylation on cell population dynamics. Specifically, it indicates a simultaneous increase in two distinct cell populations: those with bi-allelic integration and those with bi-allelic non-integration. This paradoxical outcome suggests a complex interplay between 5’ biotinylation, NHEJ inhibition, and cellular repair mechanisms.

Taken together, this study provides novel insights into the mechanisms by which 5’ modifications enhance the efficiency of precise knock-ins. These findings promote the further development of strategies for achieving high-efficiency, precise endogenous tagging in human cells.

## Acknowledgments

We thank Miho Kiyooka and Wei Chen at National Institute of Genetics for supporting PacBio sequencing, Dr. Yusuke Kishi at Institute for Quantitative Biosciences at the University of Tokyo for supporting quality control of PacBio library preparation, Dr. Mariya Genova for the proofreading of the manuscript, and the Kitagawa lab members for technical supports and helpful discussions.

## Author contributions

S.H. conceived and designed the study. R.T. designed and performed most of the experiments with help of C.T. R.T., C.T. A.M. and R.A. analyzed the PacBio data with *knock-knock*. M.F., S.Y. and T.C. provided suggestions. A.T. performed PacBio sequencing. R.T., S.H., C.T. A.M. and D.K. analyzed the data and wrote the manuscript. All authors contributed to discussions and manuscript preparation.

## Material and methods

### Cell culture

RPE1 cells, sourced from the American Type Culture Collection (ATCC), were cultured in Dulbecco’s Modified Eagle Medium/Nutrient Mixture F-12 (DMEM/F-12, Nacalai Tesque) with 10% fetal bovine serum (FBS), 100 U/mL penicillin, and 100 µg/mL streptomycin. The cells were maintained at 37°C in a humidified incubator with 5% CO2.

### Chemicals

Alt-R HDR enhancer V2 (Integrated DNA Technologies) was used as NHEJ inhibitor (NHEJi) at a concentration of 1 μM and stored at -20°C following the manufacturer’s instructions.

### Double-stranded DNA donor and guide RNA preparation

Double-stranded DNA (dsDNA) donors with or without modifications and guide RNAs (sgRNA for Cas9 and crRNA for Cpf1) were prepared according to methods described by Komori et al. (2023) and Mabuchi et al. (2023). For C-terminal and N-terminal tagging with mNeonGreen, dsDNA donors were amplified by PCR from plasmids containing the fluorescent tag and 5xGA linker sequence, flanked by 90-base homology arms (HAs). Unmodified or 5’ modified (Biotin or AmC6) primers recognizing the HA sequences were used. Unmodified dsDNA donors for mScarlet tagging were amplified by PCR from plasmids containing the mScarlet and 5xGA linker sequence. Unmodified primers containing either 90-base left or right homology arms recognizing the mScarlet or the 5xGA linker sequences were used. Modified dsDNA donors for mScarlet tagging were amplified by PCR using the unmodified mScarlet dsDNA donors as a template with 5’ modified primers recognizing the homology arms. For all PCR-amplified dsDNA donors, DpnI and exonuclease I were directly added to PCR solution to digest residual template plasmids and primers, respectively. The dsDNA donors were purified using the NucleoSpin Gel and PCR Clean-up kit (Macherey-Nagel), and then either stored at -20°C or used directly for electroporation. All primers used in this study were purchased from Eurofins Genomics and their sequences are listed in Supplementary Table 1.

Guide RNAs were synthesized from PCR-generated DNA templates. First, a DNA template containing the T7 promoter and sgRNA sequence was produced by PCR. The purified template was then transcribed *in vitro* using T7 RNA polymerase. In vitro transcription was carried out with the HiScribe T7 High Yield RNA Synthesis Kit (New England Biolabs), followed by the template DNA digestion with DNase I (Takara Bio). The synthesized guide RNA was purified using the RNA Clean & Concentrator Kit (Zymo Research). All target site sequences of the guide RNAs used in this study are listed in Supplementary Table 2.

### Doggy bone DNA donor production

Doggy bone DNA (dbDNA) donors, unique linear dsDNA structures with covalently closed ends formed by short hairpin loops, were produced through enzymatic reactions using plasmid vectors as templates. The pcDNA3.4 vectors were constructed to include two TelRL sites (TelN protelomerase recognition sequences) flanking the donor DNA sequence consists of mNeonGreen, 5xGA and HA sequences. Additionally, the backbone sequence of these template vectors contained recognition sites for NotI and XhoI, which were not included within the donor sequences. To produce dbDNA with covalentely closed ends, TelN protelomerase (0.17 U/µl, New England Biolands) was mixed with the purified plasmid vector (30 fmol/µl) in ThermoPol reaction buffer and incubated at 30°C for 30 min, followed by heat inactivation at 75 °C for 5 min. In this reaction, two types of dbDNA are produced: the desired dbDNA donors and the covalently closed backbone sequences. For the purification of dbDNA donors, unreacted template plasmids and residual backbone sequences were first digested using restriction enzymes to generate exonuclease-sensitive open ends, followed by treatment with exonucleases that are unable to cleave the covalently closed ends of dbDNA donors. After TelN protelomerase reaction, restriction enzymes NotI-HF and XhoI (both 0.4 U/µl, New England Biolands) and rCutSmart buffer were added directly to the reaction mix, incubated at 37°C for over 3 hours, followed by heat inactivation at 80°C for 20min. Exonuclease I (0.15 U/µl, New England Biolabs) and exonuclease III (0.77 U/µl, New England Biolabs) were then added with NEBuffer I to digest DNA with exposed end. The reaction mix was incubated at 37°C for 30 to 60 min, followed by heat inactivation at 80°C for 20min. Enzyme-resistant dbDNA was then column-purified using the NucleoSpin Gel and PCR Clean-up kit (Macherey-Nagel). When 18 mol of plasmid template is reacted and subsequently eluted in 30 µL of sterile MQ, 13-15 mol of dbDNA is purified, resulting in a yield of approximately 70-85%.

### Gene knock-in using the CRISPR/Cas9 and CRISPR/Cpf1 system

For both Cas9 and Cpf1-mediated endogenous tagging, gene knock-in was achieved via electroporation of RNP complexes and various DNA donor templates (unmodified, 5’ Biotin, 5’ AmC6, or dbDNA) using the Neon Transfection System (Thermo Fisher Scientific) following the manufacturer’s guidelines. HiFi Cas9 protein (1.55 µM, IDT) and sgRNA (1.84 µM) for the Cas9 system, and A.s. Cas12a Ultra (1 µM, IDT) and crRNA (1 µM) for the Cpf1 system were pre-incubated at room temperature in resuspension buffer R (Thermo Fisher Scientific). Cells (0.125 × 10^5 /µL), electroporation enhancer (Cas9 electroporation enhancer for Cas9 system, Cpf1 electroporation enhancer for Cpf1 system, 1.8 µM, IDT), and DNA donors (33 nM in total, i.e., 16.5 nM for each DNA donor in dual-color tagging) were mixed with the RNP complex for electroporation using a 10 µL Neon tip at 1300 V with two 20 ms pulses. Transfected cells were subsequently seeded into 24-well plates containing medium supplemented with DMSO or NHEJi. After 24 hours, the culture medium was replaced three times to remove the chemials.

### Fluorescent protein imaging

Five days after electroporation, cells cultured on coverslips (Matsunami) were fixed with 4% paraformaldehyde (PFA) at room temperature for 15 minutes and subsequently washed with PBS. Permeabilization was then achieved by treating the cells with 0.05% Triton X-100 in PBS at room temperature for 10 minutes, followed by PBS wash. The cells were then mounted onto glass slides (Matsunami) using ProLong Gold Antifade Mountant with DAPI (Invitrogen). Images were acquired using an Axio Imager.M2 microscope (Carl Zeiss) equipped with a 63× lens objective.

### Quantification of knock-in efficiency by flow cytometry

4 days post-electroporation, cells were subjected to flow cytometric analysis. The adherent cells were detached using a trypsin/EDTA solution and resuspended in PBS. A BD FACS Aria III system (BD Biosciences), equipped with 355/405/488/561/633 nm lasers, was employed to detect mNG-positive cells in the suspension. Data were collected from more than 10,000 gated events.

### Amplicon sequencing and analysis by *knock-knock*

#### Genomic DNA preparation

Amplicon sequencing and subsequent analysis using the *knock-knock* platform were conducted following methods described in Tei et al. (2023). For unsorted samples, cells were maintained in media containing DMSO or NHEJi for 24 h after electroporation, then cultured for an additional 3 days as described above. Genomic DNA was then isolated using NucleoSpin DNA RapidLyse kit (Macherey-Nagel). For fluorescence-sorted cells, after electroporation and subsequent cell culture, a minimum of 6,200 cells exhibiting mNG fluorescence were isolated using a FACS Aria III system. The genomic DNA from these sorted cells was then extracted using the NucleoSpin Tissue XS kit (Macherey-Nagel).

#### Amplicon sequencing

Amplicon libraries were prepared using a two-step PCR and adapter ligation method following the instructions by Pacific Biosciences (Part Number 101-791-800 Version 02, April 2020) with minor adjustments. In the initial PCR amplification, extracted genomic DNA was used as a template to amplify a segment flanking the target site of mNG insertion. The amplification was performed using KOD One Master Mix (TOYOBO) and primers with universal sequences which serve as annealing sites for barcoded primers in subsequent reactions. Following purification with AMPure XP beads (Beckman Coulter), the amplified DNA underwent a second round of PCR using primers sourced from the Barcoded Universal F/R Primers Plate-96v2 (Pacific Biosciences). The resulting products were subsequently purified using AMPure PB beads (Pacific Biosciences). The barcoded amplicons were analyzed using TapeStation (Agilent Technologies) and Qubit Fluorometer (Thermo Fisher Scientific). Finally, all the amplicons were pooled together in equimolar amounts as a single sample. The amplicon pooled libraries were prepared using the SMRTbell Express Template Prep Kit 2.0 (Pacific Biosciences, CA, USA). Four Sequel II SMRT Cells were run on the PacBio Sequel II/IIe systems with Binding Kit 3.1 or 3.2/Sequencing Kit 2.0 (Pacific Biosciences, CA, USA) and 30 hour movies. The consensus (HiFi) reads were generated from the raw full-pass subreads using the DeepConsensus v1.2 program (Baid et al., 2023), and then were demultiplexed by sample barcodes using the SMRT Link v13.1.0.221970 software (Pacific Biosciences, CA, USA). A total of 2,488,194 barcoded Reads with read quality score >= 40 were selected.

#### Analysis of knock-in outcomes by knock-knock

For the analysis of knock-in outcomes using *knock-knock*, a computational pipeline developed by Canaj et al. (2019), all barcoded reads underwent trimming of universal sequences. The knock-knock software package utilized in our study is available at https://github.com/jeffhussmann/knock-knock. Based on the detailed classification provided by *knock-knock*, repair patterns that were classified as “simple indel”, “complex indel” and “genomic insertion(hg38)” were grouped as “indels”. These “indels” along with “WT” were collectively categorized as “donor-independent repair”. Other repair patterns, classified as “HDR”, “blunt mis-integration”, “incomplete HDR”, “donor fragment”, “complex mis-integration” and “concatenated mis-integration”, were grouped as “donor-dependent repair”. For the detailed categorization of donor-dependent repair, in each figure, repair patterns that were subcategorized as “5’ blunt, 3’ blunt” by *knock-knock* classification were grouped as “blunt”. Repair patterns subcategorized as “5’ HDR, 3’ blunt”, “5’ blunt, 3’ HDR”, “5’ HDR, 3’ imperfect”, and “5’ imperfect, 3’ HDR” were grouped as “asymmetric HDR”. Repair patterns subcategorized as “5’ blunt, 3’ imperfect”, “5’ imperfect, 3’ blunt”, and “5’ imperfect, 3’ imperfect” were grouped to “imperfect”. For the detailed categorization of donor independent repair, repair patterns that were classified as “genomic insertion(hg38)” were included in “complex indel” group in supplementary Figure.

#### Statistical analysis

Statistical comparisons between data from different groups were conducted using PRISM v.10 software (GraphPad). Depending on the context, either a Tukey–Kramer test or a two-tailed, unpaired Student’s t-test was employed, as specified in the figure legends. P values less than 0.05 were considered statistically significant. All data shown are mean ± S.D., with sample sizes indicated in the figure legends.

---

**Fig. S1 related to Fig. 1**

**a**, Representative images from cells with Cpf1-mediated mNG tagging of CLTC. Arrowheads indicate cells with cytoplasmic mNG signals. Cells at 5 days after electroporation were fixed and analyzed. Scale bar: 10 µm. **b**, Percentage of WT and indels within donor-independent repair across the targeted genes. For each sample, 9,386-45,809 reads categorized as the donor-independent repair were analyzed. Data are represented as mean ± S.D.

**Fig. S2 related to Fig. 2**

**a**, Percentage of WT and indels within donor-independent repair across the targeted genes in Fig. 2d. For each sample, 10,952-146,558 reads categorized as the donor-independent repair were analyzed. Data are represented as mean ± S.D.

**Fig. S3 related to Fig. 3**

**a**, Percentage of WT and indels within donor-independent repair across the targeted genes in Fig. 3c. For each sample, 7,176-45,809 reads categorized as the donor-independent repair were analyzed. Data are represented as mean ± S.D.

**Fig. S4 related to Fig. 4**

**a**, Percentage of WT and indels within donor-independent repair across the targeted genes in Fig. 4a. For each sample, 4,610-25,938 reads categorized as the donor-independent repair were analyzed. Data are represented as mean ± S.D.

## References

Allen, A., Wang, C., Caproni, L. J., Sugiyarto, G., Harden, E., Douglas, L. R., Duriez, P. J., Karbowniczek, K., Extance, J., Rothwell, P. J., et al. (2018). Linear doggybone DNA vaccine induces similar immunological responses to conventional plasmid DNA independently of immune recognition by TLR9 in a pre-clinical model. *Cancer Immunology*, Immunotherapy 67, 627–638.

Arturo Gutierrez-Triana, J., Tavhelidse, T., Thumberger, T., Thomas, I., Wittbrodt, B., Kellner, T., Anlas, K., Tsingos, E. and Wittbrodt, J. (2018). Efficient single-copy HDR by 5’ modified long dsDNA donors. eLife 7, e39468.

Baid, G., Cook, D. E., Shafin, K., Yun, T., Llinares-López, F., Berthet, Q., Belyaeva, A., Töpfer, A., Wenger, A. M., Rowell, W. J., et al. (2023). DeepConsensus improves the accuracy of sequences with a gap-aware sequence transformer. Nat Biotechnol 41, 232–238.

Bhargava, R., Onyango, D. O. and Stark, J. M. (2016). Regulation of Single-Strand Annealing and its Role in Genome Maintenance. Trends in Genetics 32, 566– 575.

Bishop, D. C., Caproni, L., Gowrishankar, K., Legiewicz, M., Karbowniczek, K., Tite, J., Gottlieb, D. J. and Micklethwaite, K. P. (2020). CAR T Cell Generation by piggyBac Transposition from Linear Doggybone DNA Vectors Requires Transposon DNA-Flanking Regions. Mol Ther Methods Clin Dev 17, 359–368.

Canaj, H., Hussmann, J. A., Li, H., Beckman, K. A., Goodrich, L., Cho, N. H., Li, Y. J., Santos, D. A., Mcgeever, A., Stewart, E. M., et al. (2019). Deep profiling reveals substantial heterogeneity of integration outcomes in CRISPR knock-in experiments. BioRxiv. 10.1101/841098.

Ceccaldi, R., Rondinelli, B. and D’Andrea, A. D. (2016). Repair Pathway Choices and Consequences at the Double-Strand Break. Trends Cell Biol 26, 52–64.

Chang, H. H. Y., Watanabe, G. and Lieber, M. R. (2015). Unifying the DNA end-processing roles of the artemis nuclease: Ku-dependentartemis resection atbluntdnaends. Journal of Biological Chemistry 290, 24036–24050.

Chang, H. H. Y., Pannunzio, N. R., Adachi, N. and Lieber, M. R. (2017). Non-homologous DNA end joining and alternative pathways to double-strand break repair. Nat Rev Mol Cell Biol 18, 495–506.

Chen, F., Pruett-Miller, S. M., Huang, Y., Gjoka, M., Duda, K., Taunton, J., Collingwood, T. N., Frodin, M. and Davis, G. D. (2011). High-frequency genome editing using ssDNA oligonucleotides with zinc-finger nucleases. Nat Methods 8, 753–757.

Decottignies, A. (2013). Alternative end-joining mechanisms: A historical perspective. Front Genet 4, 1–7.

Deneke, J., Nter Ziegelin, G., Lurz, R. and Lanka, E. (2002). Phage N15 Telomere Resolution TARGET REQUIREMENTS FOR RECOGNITION AND PROCESSING BY THE PROTELOMERASE*. JBC Papers in Press 277, 10410–10419.

Denes, C. E., Cole, A. J., Aksoy, Y. A., Li, G., Neely, G. G. and Hesselson, D. (2021). Approaches to enhance precise crispr/cas9-mediated genome editing. Int J Mol Sci 22, 8751.

Elda Cannavo & Petr Cejka. (2014). Sae2 promotes dsDNA endonuclease activity within Mre11–Rad50–Xrs2 to resect DNA breaks. Nature 514, 7520.

Feng, S., Sekine, S., Pessino, V., Li, H., Leonetti, M. D. and Huang, B. (2017). Improved split fluorescent proteins for endogenous protein labeling. Nat Commun 8, 370.

Fu, Y. W., Dai, X. Y., Wang, W. T., Yang, Z. X., Zhao, J. J., Zhang, J. P., Wen, W., Zhang, F., Oberg, K. C., Zhang, L. (2021). Dynamics and competition of CRISPR-Cas9 ribonucleoproteins and AAV donor-mediated NHEJ, MMEJ and HDR editing. Nucleic Acids Res 49, 969–985.

Ghanta, K. S., Chen, Z., Mir, A., Dokshin, G. A., Krishnamurthy, P. M., Yoon, Y., Gallant, J., Xu, P., Zhang, X.-O., Rasit Ozturk, A. (2021). 5′-Modifications improve potency and efficacy of DNA donors for precision genome editing. eLife 10, e72216.

Heinrich, J., Schultz, J., Bosse, M., Ziegelin, G., Lanka, E. and Moelling, K. (2002). Linear closed mini DNA generated by the prokaryotic cleaving-joining enzyme TelN is functional in mammalian cells. J Mol Med 80, 648–654.

Hsu, P. D., Lander, E. S. and Zhang, F. (2014). Development and applications of CRISPR-Cas9 for genome engineering. Cell 157, 1262–1278.

Jinek, M., Chylinski, K., Fonfara, I., Hauer, M., Doudna, J. A. and Charpentier, E. (2012). A Programmable Dual-RNA–Guided DNA Endonuclease in Adaptive Bacterial Immunity. Science (1979) 337, 816–821.

Karda, R., Counsell, J. R., Karbowniczek, K., Caproni, L. J., Tite, J. P. and Waddington, S. N. (2019). Production of lentiviral vectors using novel, enzymatically produced, linear DNA. Gene Ther 26, 86–92.

Lovett, S. T. (2011). The DNA Exonucleases of Escherichia coli. EcoSal Plus 4,.

Mabuchi, A., Hata, S., Genova, M., Tei, C., Ito, K. K., Hirota, M., Komori, T., Fukuyama, M., Chinen, T., Toyoda, A., et al. (2023). ssDNA is not superior to dsDNA as long HDR donors for CRISPR-mediated endogenous gene tagging in human diploid RPE1 and HCT116 cells. BMC Genomics 24, 289.

Maruyama, T., Dougan, S. K., Truttmann, M. C., Bilate, A. M., Ingram, J. R. and Ploegh, H. L. (2015). Increasing the efficiency of precise genome editing with CRISPR-Cas9 by inhibition of nonhomologous end joining. Nat Biotechnol 33, 538–542.

Sayers, J. R. and Ecksteins, F. (1990). Properties of Overexpressed Phage T5 D15 Exonuclease SIMILARITIES WITH ESCHERICHIA COLI DNA POLYMERASE 15’-3’ EXONUCLEASE*. JCB 265, 30

Schimmel, J., Muñoz-Subirana, N., Kool, H., van Schendel, R., van der Vlies, S., Kamp, J. A., de Vrij, F., Kushner, S. A., Smith, G. C. M., Boulton, S. J., et al. (2023). Modulating mutational outcomes and improving precise gene editing at CRISPR-Cas9-induced breaks by chemical inhibition of end-joining pathways. Cell Rep 42, 112019.

Scully, R., Panday, A., Elango, R. and Willis, N. A. (2019). DNA double-strand break repair-pathway choice in somatic mammalian cells. Nat Rev Mol Cell Biol 20, 698–714.

Seol, J. H., Shim, E. Y. and Lee, S. E. (2018). Microhomology-mediated end joining: Good, bad and ugly. Mutation Research - Fundamental and Molecular Mechanisms of Mutagenesis 809, 81–87.

Sfeir, A. and Symington, L. S. (2015). Microhomology-Mediated End Joining: A Back-up Survival Mechanism or Dedicated Pathway? Trends Biochem Sci 40, 701–714.

Sinha, S., Villarreal, D., Shim, E. Y. and Lee, S. E. (2016). Risky business: Microhomology-mediated end joining. Mutation Research - Fundamental and Molecular Mechanisms of Mutagenesis 788, 17–24.

Tei, C., Hata, S., Mabuchi, A., Okuda, S., Ito, K. K., Genova, M., Fukuyama, M., Yamamoto, S., Chinen, T., Toyoda, A., et al. (2023). Comparable analysis of multiple DNA double-strand break repair pathways in CRISPR-mediated endogenous tagging. BioRxiv. 10.1101/2023.06.28.546861.

Thornton, C. D., Fielding, S., Karbowniczek, K., Roig-Merino, A., Burrows, A. E., FitzPatrick, L. M., Sharaireh, A., Tite, J. P., Mole, S. E., Harbottle, R. P., et al. (2021). Safe and stable generation of induced pluripotent stem cells using doggybone DNA vectors. Mol Ther Methods Clin Dev 23, 348–358.

van de Kooij, B., Kruswick, A., van Attikum, H. and Yaffe, M. B. (2022). Multi-pathway DNA-repair reporters reveal competition between end-joining, single-strand annealing and homologous recombination at Cas9-induced DNA double-strand breaks. Nat Commun 13, 5295.

Wu, Y., Liang, D., Wang, Y., Bai, M., Tang, W., Bao, S., Yan, Z., Li, D. and Li, J. (2013). Correction of a genetic disease in mouse via use of CRISPR-Cas9. Cell Stem Cell 13, 659–662.

Yao, R., Liu, D., Jia, X., Zheng, Y., Liu, W. and Xiao, Y. (2018). CRISPR-Cas9/Cas12a biotechnology and application in bacteria. Synth Syst Biotechnol 3, 135–149.

Yu, C., Liu, Y., Ma, T., Liu, K., Xu, S., Zhang, Y., Liu, H., La Russa, M., Xie, M., Ding, S., et al. (2015). Small molecules enhance crispr genome editing in pluripotent stem cells. Cell Stem Cell 16, 142–147.

Yu, Y., Guo, Y., Tian, Q., Lan, Y., Yeh, H., Zhang, M., Tasan, I., Jain, S. and Zhao, H. (2020). An efficient gene knock-in strategy using 5′-modified double-stranded DNA donors with short homology arms. Nat Chem Biol 16, 387–390.

Zetsche, B., Gootenberg, J. S., Abudayyeh, O. O., Slaymaker, I. M., Makarova, K. S., Essletzbichler, P., Volz, S. E., Joung, J., Van Der Oost, J., Regev, A., et al. (2015). Cpf1 Is a Single RNA-Guided Endonuclease of a Class 2 CRISPR-Cas System. Cell 163, 759–771.

Zhang, X., Li, T., Ou, J., Huang, J. and Liang, P. (2022). Homology-based repair induced by CRISPR-Cas nucleases in mammalian embryo genome editing. Protein Cell 13, 316–335.

Zhao, Z., Shang, P., Sage, F. and Geijsen, N. (2022). Ligation-assisted homologous recombination enables precise genome editing by deploying both MMEJ and HDR. Nucleic Acids Res 50, E62.

