## Supplementary figures and images for "Comprehensive analysis of end-modified long dsDNA donors in CRISPR-mediated endogenous tagging"

### supplementary figure

#### Supplementary Figure 1

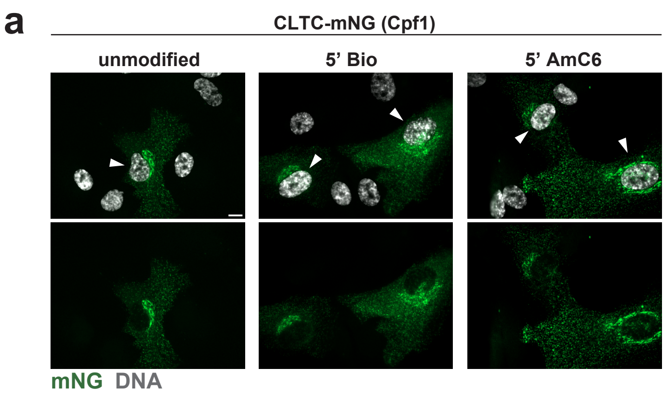


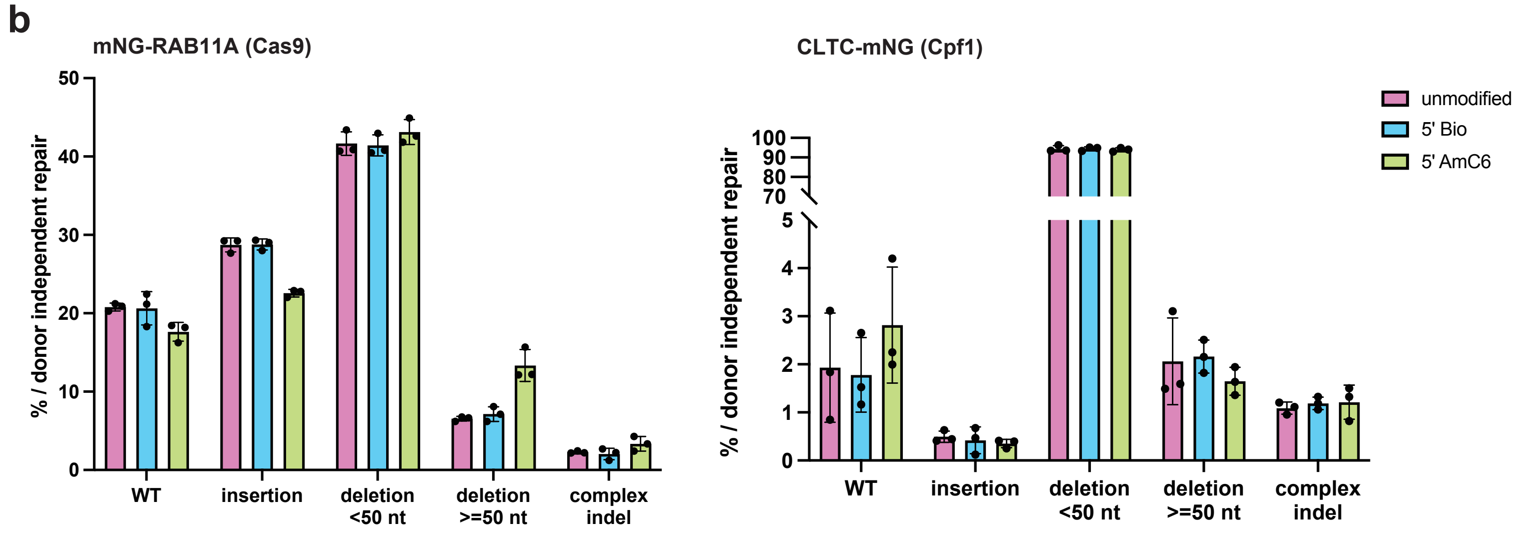

#### Supplementary Figure 2

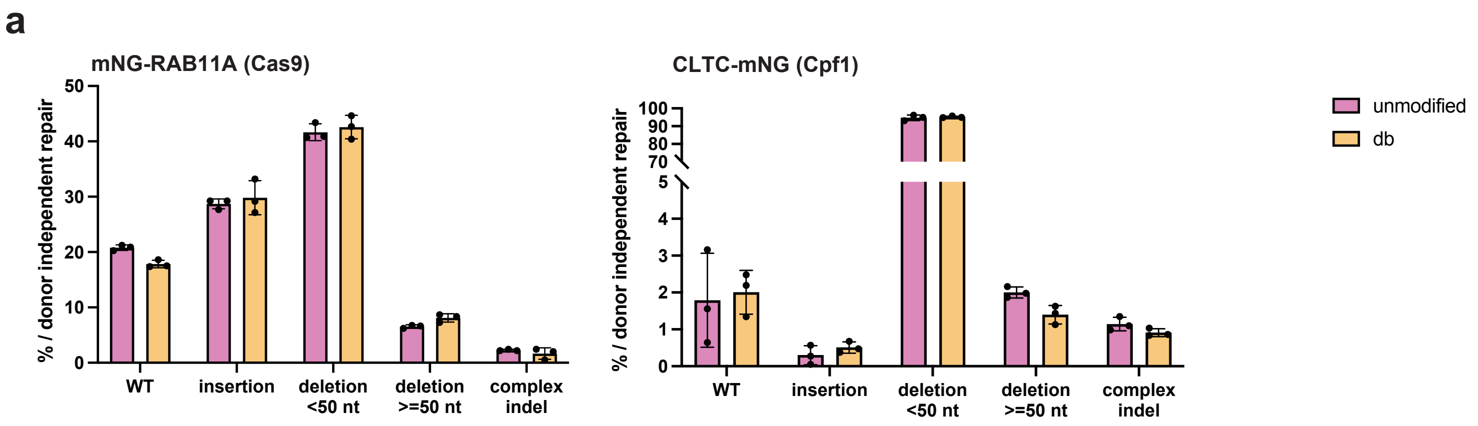

#### Supplementary Figure 3

**a**

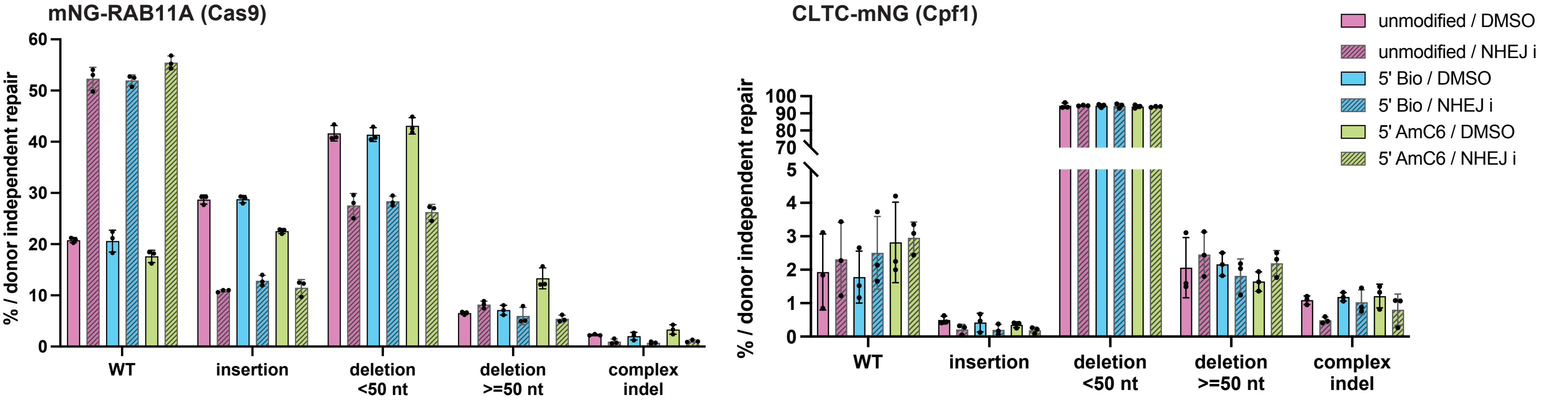

#### Supplementary Figure 4

**a**

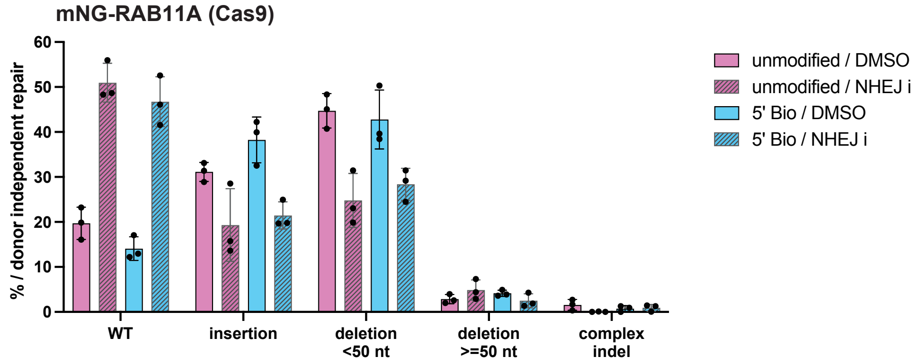
