## supplementary table for "Comprehensive analysis of end-modified long dsDNA donors in CRISPR-mediated endogenous tagging"

### Supplementary Table 1: Primer sequences for PCR

#### For guide RNA assembly

| Name | Sequence |
|---|---|
| crRNA_tracrRNA | GTTTTAGAGCTAGAAATAGCAAGTTAAAATAAGGCTAGTCCGTTATCAACTTGAAA AAGTGGCACCGAGTCGGTGCTTTT |
| RAB11A_sgRNA_Fw | TTCTAATACGACTCACTATAGGTAGTCGTACTCGTCGTCG |
| RAB11A_sgRNA_ Rv | TTCTAGCTCTAAAACCGACGACGAGTACGACTACC |
| Univ_sgRNA_Fw | TTCTAATACGACTCACTATAG |
| Univ_sgRNA_Rv | AAAAGCACCGACTCGGTG |
| Cpf1_crRNA_Fw | TTCTAATACGACTCACTATAGTAATTTCTACTCTTGTAGAT |
| CLTC_sgRNA_Rv | TTCATCTCACATGCTGTACCATCTACAAGAGTAGAAATTAC |

#### For HDR donor preparation

| Name | Sequence |
|---|---|
| mNG-RAB11A(unmodified, 5' Bio, 5' AmC6)<br>mSc-RAB11A(5' Bio, 5' AmC6)_Fw | GGCGCTCGGGTTACCCCTG |
| mNG-RAB11A(unmodified, 5' Bio, 5' AmC6)<br>mSc-RAB11A(5' Bio, 5' AmC6)_Rv | GGGAGTGGCCCGGGTCCC |
| mSc-RAB11A(unmodified)_Fw | GGCGCTCGGGTTACCCCTGCAGCGACGCCCCCTGGTCCCACAGATACCACTGCTGCTCCCGCCCTTTCGCTCCTCGGCCGCGCAATGGGCATGGTGAGCAAGGGCG AGGC |
| mSc-RAB11A(unmodified)_Rv | GGGAGTGGCCCGGGTCCCCGAACGAGGACTGTGTAGAGTGCGAGAGCCCATGGCCTCACCTTTAAAGAGGTAGTCGTACTCGTCGTCGCGTGCACCAGCTCCTGCA CC |
| CLTC-mNG(unmodified, 5' Bio, 5' AmC6)<br>CLTC-mSc(5' Bio, 5' AmC6)_Fw | AGTGTTGCCGTCCCTCCC |
| CLTC-mNG(unmodified, 5' Bio, 5' AmC6)<br>CLTC-mSc(5' Bio, 5' AmC6)_Rv | GTTGCCTGTTTTCCCCCATTATAAAC |
| CLTC-mSc(unmodified)_Fw | AGTGTTGCCGTCCCTCCCCAGGCACCTTTTGGTTATGGTTATACCGCACCACCGTATGGACAGCCACAGCCTGGCTTTGGGTACAGCATGGGAGCTGGTGCAGGTGCAG |
| CLTC-mSc(unmodified)_Rv | GTTGCCTGTTTTCCCCCATTATAAACTGAGAAGTGGGTAAAGACGATGTTTCAGTACGAAAATAGGTGACTACAGGATCAGCGCTTCATCCTACTTGTACAGCTCGTCCATGC |

**For long-read amplicon sequencing**

| Name | Sequence |
|---|---|
| SMRT_1st_Fw_RAB11A | [AmC6]GCAGTCGAACATGTAGCTGACTCAGGTCACAGTGAGTGACAAGCGCCTAC |
| SMRT_1st_Rv_RAB11A | [AmC6]TGGATCACTTGTGCAAGCATCACATCGTAGCTCAGGAACCAGGCCTTCAG |
| SMRT_1st_Fw_CLTC | [[AmC6]GCAGTCGAACATGTAGCTGACTCAGGTCACTTGGCTTGCCTTCAGGGTGTTTTC |
| SMRT_1st_Rv_CLTC | [AmC6]TGGATCACTTGTGCAAGCATCACATCGTAGGCCTCCCTAATGCCTCAGTATCCA |

### Supplementary Table 2: Target site sequences of guide RNA

| Target gene | Cas nuclease | Target sequence |
|---|---|---|
| *RAB11A* (Human) | Cas9 | GGTAGTCGTACTCGTCGTCG |
| *CLTC* (Human) | Cpf1 | GGTACAGCATGTGAGATGAA |
